# Reproducible Tract Profiles (RTP): from diffusion MRI acquisition to publication

**DOI:** 10.1101/680173

**Authors:** Garikoitz Lerma-Usabiaga, Michael L. Perry, Brian A. Wandell

**Affiliations:** Department of Psychology, Stanford University, 450 Serra Mall, Jordan Hall Building, 94305 Stanford, California, USA; BCBL. Basque Center on Cognition, Brain and Language. Mikeletegi Pasealekua 69, Donostia - San Sebastián, 20009 Gipuzkoa, Spain

**Author notes:** **Corresponding Author:** Garikoitz Lerma-Usabiaga, 450 Serra Mall, Room 488, Jordan Hall, Building 420, Main Quad, 94305 Stanford CA, United States.

**Keywords:** Computational Reproducibility, Diffusion MRI, DWI, White Matter Tracts

## Abstract

Reproducible Tract Profiles (RTP) comprises a set of methods to manage and analyze diffusion weighted imaging (DWI) data for reproducible tractography. The tools take MRI data from the scanner and process them through a series of analysis implemented as Docker containers that are integrated into a modern neuroinformatics platform (Flywheel). The platform guarantees that the entire pipeline can be re-executed, using the same data and computational parameters. In this paper, we describe (1) a cloud based neuroinformatics platform, (2) a tool to programmatically access and control the platform from a client, and (3) the DWI analysis tools that are used to identify the positions of 22 tracts and their diffusion profiles. The combination of these three components defines a system that transforms raw data into reproducible tract profiles for publication.

**Graphical abstract:** Reproducible Tract Profiles (RTP) comprises a set of methods to manage and analyze diffusion weighted imaging (DWI) data for reproducible tractography. The RTP methods comprise two main parts.

1. Server side software tools for storing data and metadata and managing containerized computations.
2. Client side software tools that enable the researcher to read data and metadata and manage server-side computations.

The server-side computational tools are embedded in containers that are linked to a JSON file with a complete specification of the computational parameters. The data and computational infrastructure on the server is fully reproducible.

## Introduction

The increasing complexity of neuroimaging data and analyses is a complicating factor that challenges researchers’ ability to implement reproducible research methods. In many neuroimaging publications, there is no realistic chance that a reader can replicate the experimental data acquisition or reproduce the computational analyses (Wilson et al. 2017; Sandve et al. 2013; Buckheit and Donoho 1995). It is understandable that the costs of replicating an experiment, particularly with a special patient cohort, may be impractical. It should still be possible to share the data and enable colleagues to repeat, check, and explore critical portions of the computational analysis in a publication (Stodden, Leisch, and Peng 2014; Peng 2011).

At this moment, the increase in computational power has mainly led to an increase in algorithm complexity, and the associated parameters. Choices made in these pipelines can have substantial effects on the reported results. In the case of functional MRI (fMRI), the position of the peak activation may range over a cortical area of 25 cm^2^ (Carp 2012). Further, (Yarkoni and Westfall 2017) observe that we are often uncertain about critical parameters that must be in computational models. We confirm that this observation also applies to DWI and tractography. It is our experience, too, that scientists find it very difficult to keep track of the full range of parameters used in any particular analysis, and even fewer scientists record the combinations of parameters they used during data exploration (Baker 2016).

To address this problem, we are developing tools that use the increased computational power to improve computational reproducibility. In this paper we describe methods that implement Reproducible Tract Profiles (RTP). These methods implement fully reproducible analyses that begin with data acquisition and end with the final figures used in a publication. The RTP methods include the computational steps required for a complete DWI analysis solution. This solution is reproducible because the analysis software and its dependencies are encapsulated in a Docker container, and the input data, complete set of analysis configuration parameters and the outputs are stored in a single neuroinformatics platform (Flywheel). The neuroinformatics platform runs on a server in the cloud. The user can access the data and control the computations from the client side using another provided tool (Scitran), that programmatically controls the interactions with the platform. Together the RTP tools comprise a system that enables users to check and reproduce all the computations from DWI acquisition to publication.

## Method description

We divided the description of the methods into two parts: (1) *the infrastructure:* required for a computationally reproducible system, sometimes called the neuroinformatics platform (Marcus et al. 2011); and, (2) *the DWI data analysis pipeline*: comprising all the steps starting with the DICOM images generated in the MRI scanner (the acquisition device) to the final published results. In Figure 1, we would consider the neuroinformatics platform to be everything except what is being run inside the Docker containers (Gears, in Flywheel terminology). The data analysis pipeline is the set of operations performed to the MRI data inside the Gears.

**Figure 1.**
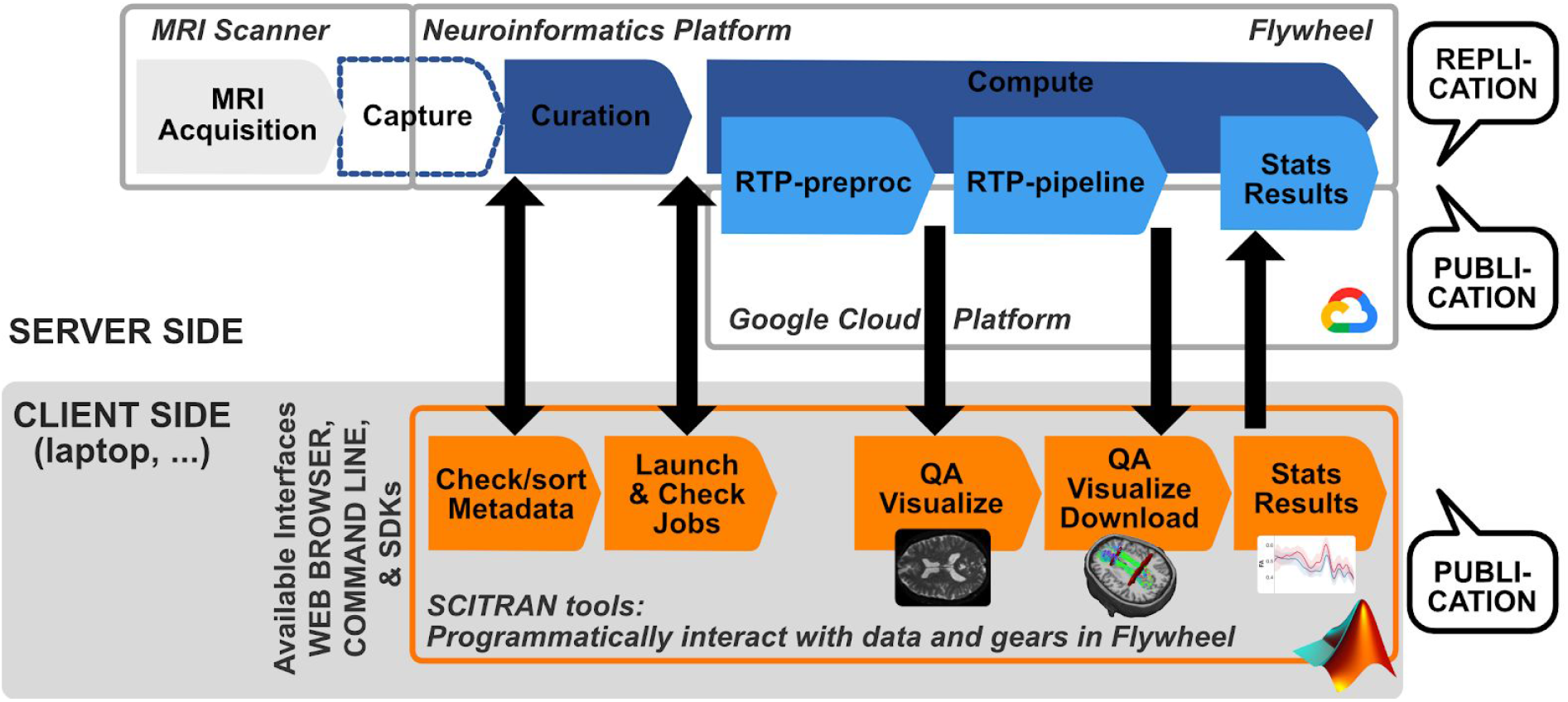
Summary of the Reproducible Tract Profiles system and associated infrastructure. The images acquired from the MRI scanner are automatically transferred to the neuroinformatics platform (Flywheel). Docker images containing the DWI analysis tools along with files specifying the computational parameters are stored and managed in the Flywheel system. This server-side system runs on the Google Cloud Platform. Data and computations in Flywheel can be read and managed programmatically from the client using a set of Matlab methods, called Scitran. These methods enable the researcher to control Flywheel data and computations, including data/metadata management, launching and checking computations, and visualizing data and results. The data and fully computational specification are stored within Flywheel, rendering the analyses fully reproducible.

### Infrastructure for computational reproducibility

#### Functionality

The data management and computational infrastructure use a technology (Flywheel.io; see Figure 2) that (a) implements reproducible computational methods, (b) tracks provenance of the data, and (c) facilitates data sharing. In Figure 2A we observe the main functionalities provided by Flywheel:

1. **Capture**: data acquired in the MRI scanner is uploaded directly to the Flywheel platform with no human interaction and based in a set of rules. Some automatic tasks are performed in this step, such as DICOM to Nifti conversion or categorization according to acquisition types. The automatic steps are configurable according to acquisition site or project specifications.
2. **Curate**: once the data has been uploaded, it becomes available for the researcher to interact with it. Data can be reorganized, shared with other project members, renamed, and most importantly, subject and data metadata can be added. These steps are not automatic, they can be done using the web interface, or more interestingly, using scripts (R, Python, Matlab) that allow changing multiple data acquisitions at once and programmatically. For this task, we provide a Matlab tool called scitran (https://github.com/vistalab/scitran/), that will be detailed below.
3. **Compute**: The neuroinformatics platform provides a system for analyzing the data in a reproducible manner. Docker containers (called Gears in Flywheel) that perform analysis are installed in the system and perform analysis over the data. Every analysis is stored independently, with the inputs and outputs and the version of the gear that was used and all the parameters.
4. **Collaborate**: All the data in the system can be easily made available to anyone in the same organization. For people outside the organization, an external user can be created for collaboration or the data and the results can be exported in BIDS or a proprietary format. To track the provenance, the computational system stores: (a) the input data, (b) the container version that was executed, (c) the container input parameters, and (c) the output files. The analyses are fully reproducible by anyone with IRB authorization to access the system. The input data, exact version of the gears and the configuration parameters can be exported as Docker images as well. These Docker images can be run in any system capable of running Docker or singularity containers. Therefore, having a Flywheel infrastructure is not necessary to be able to reproduce any of our results.

**Figure 2.**
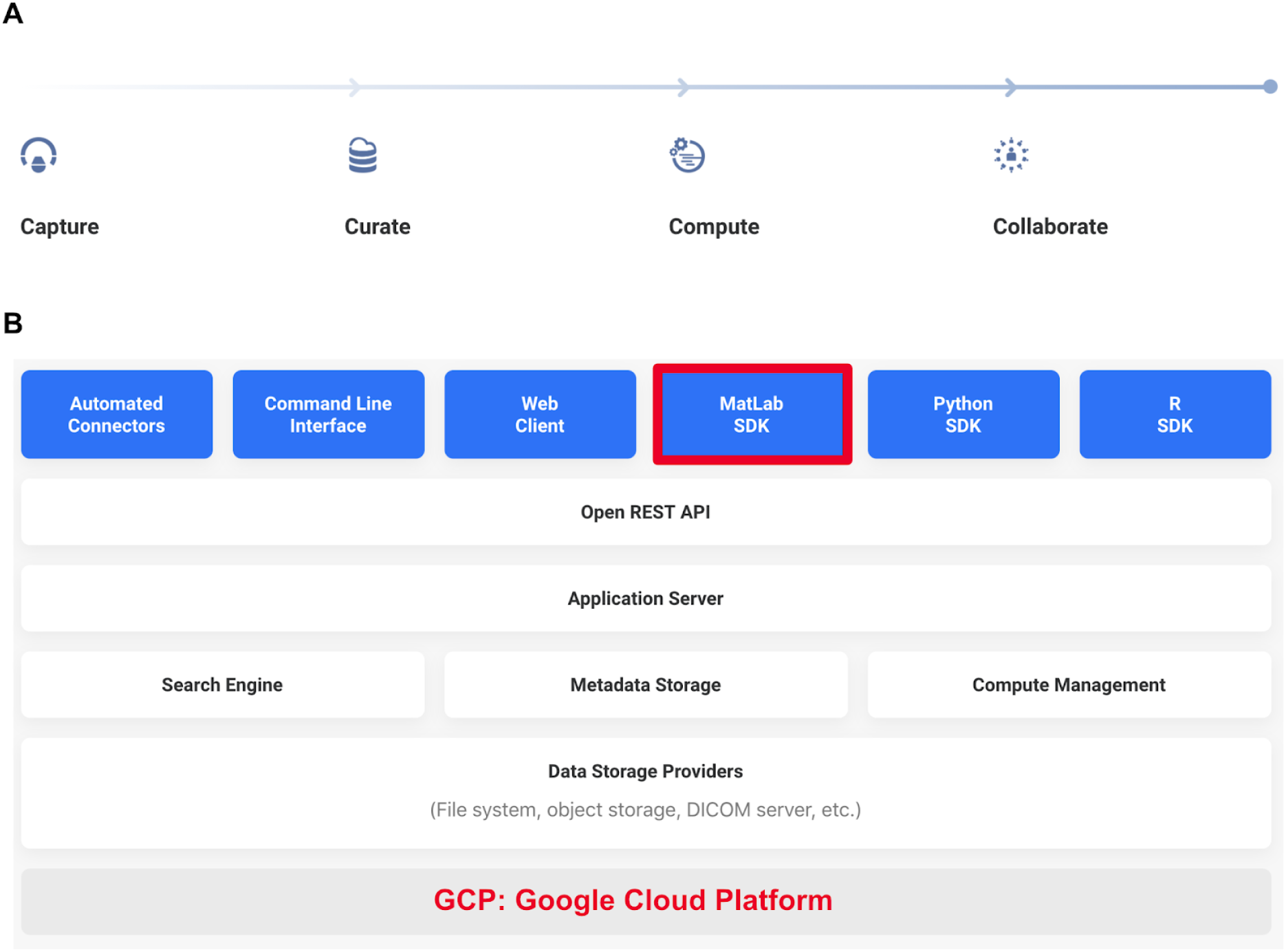
Flywheel’s main functionalities and technical architecture. A) Main functionalities covered by Flywheel’s neuroinformatics platform. B) Main components of Flywheel’s technical architecture. See main text for detailed explanations.

#### Technical architecture

Flywheel’s technical architecture is shown in Figure 2B. The first row in blue shows the different ways to interact with the system: *(1) Automated connectors:* load data from the MRI scanners to Flywheel. *(2) Command Line Interface:* a program installed locally allows the authentication to the Flywheel system and most of the management operations. Useful, for example, to upload existing BIDS formated datasets to Flywheel if we don’t have access to the original scanner. *(3) Web Client:* web interface to the Flywheel system accessible from all the major browsers. Allows the majority of the operations with a clean point and click interface. *(4-5-6) Matlab-Python-R SDK:* the SDK gives programmatic access to the Flywheel system, with the three main scientific programming languages. We marked Matlab SDK in red in Figure 2B because as part of the RTP solution we included Scitran, which is a collection of Matlab scripts designed to interact programmatically with Flywheel (see next section).

These interfaces use the Open (public) REST API (Representational State Transfer Application Programming Interface) to access Flywheel’s Application Server, which control the web interface and Flywheel’s three main systems: *(1) search engine:* accessible through the web GUI and from the SDK to search and retrieve only the required information, *(2) Metadata Storage:* core of the system, with references to all content, gear and analyses, and *(3) Compute management:* it manages the dispatching of jobs, i.e. it takes the input data, runs the Gear, and when finished, copies the results back to Flywheel.

The last two rows represent the data storage and the actual hardware where Flywheel is running (and where the Gears run as well). In our specific instance of Flywheel, both the data and the computations happen in the Google Cloud Platform. Small Gears run in the same hardware as Flywheel, but big Gears (as RTP-preproc and RTP-pipeline) launch a new virtual machine each time, runs the Gear, and shuts it off when finished. There is a parameter in Flywheel that controls the number of machines that can run in parallel. This is required for both technical and economical reasons, as Google is not usually the bottleneck in this situation.

#### Gear creation and configuration

The gear creation process starts in a client machine. After testing that our analysis tool works locally (RTP-preproc or RTP-pipeline in our case), we start the Docker Image creation process. Docker is the platform where the Docker images run, and a running Image is called a Container, i.e. a Docker Container is the run-time instance of a Docker Image. The first step is selecting a base Image that has the features we require. For example, as RTP-pipeline is a Matlab based program, we select a base Docker Image with the Matlab r2017a runtime, which is itself based on Ubuntu. In this base Docker Image, we install all the required programs, for example mrTrix (for an updated complete list of installed programs, refer to the Dockerfile file in https://github.com/vistalab/RTP-pipeline). At this point, if it is a Matlab based program, we compile it, otherwise, we build the Docker Image. Building the Image means obtaining all the required files and programs, including the compiled Matlab program if required.

Until this moment, the Gear creation was no different from a normal Docker image creation program. To make it a Flywheel Gear, we need to create a manifest.json file. The manifest.json file contains all the information relevant to the Gear, such as name, version, labels, description, inputs, outputs, and configuration parameter types and defaults. See the RTP-pipeline github page for an example manifest.json file. The manifest tells Flywheel what to expect from this Docker container, what inputs should it take, and how to construct the web client gui to be able to edit the configuration parameters. This is an example of how to set an input file in the manifest.json file:

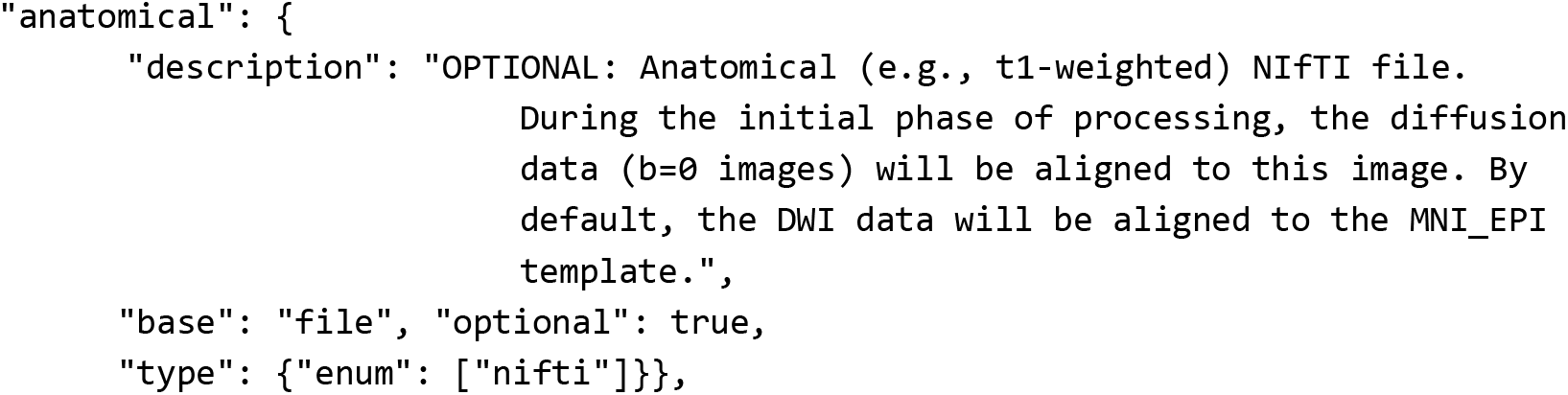

And this is an example of how to set up a config parameter:

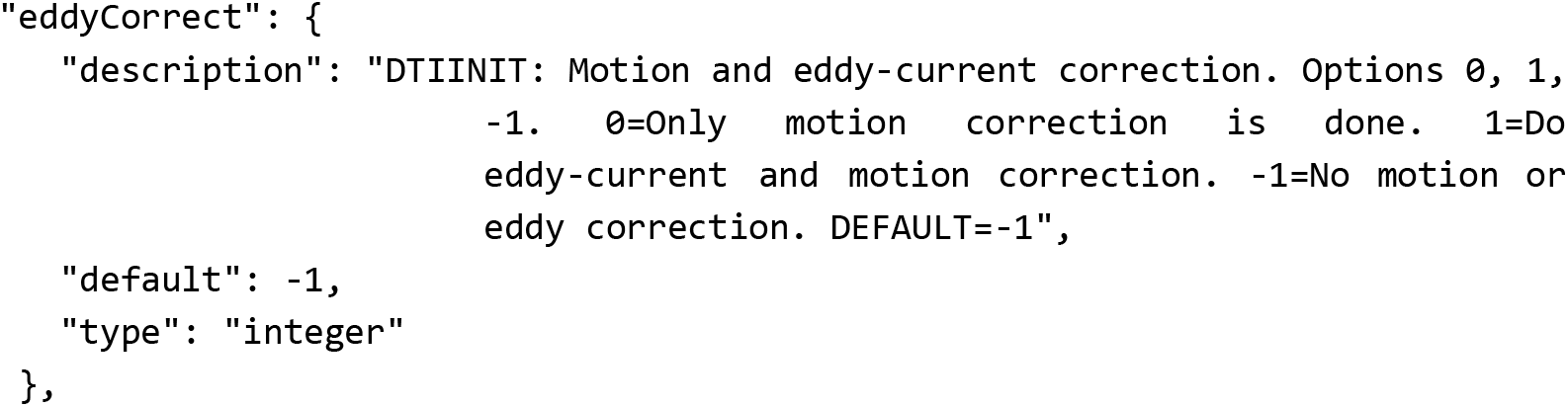

When we run the Docker container without passing configuration parameters, it will run it using the defaults from the manifest.json. Once we have the Docker Container working locally, we can upload it and install it in Flywheel. For a detailed Gear creation and installation process, please refer to Flywheel’s official documentation in https://docs.flywheel.io.

The full cycle of testing a gear has the following steps: (1) Run the code directly in Matlab and check the outputs; (2) Compile the Matlab code and run it using Matlab’s Runtime environment (it is freely downloadable from www.mathworks.com); (3) Build the Docker Container and run it using the default parameters and changed parameters; (4) Check the code in github, tag it, and build the containers in dockerhub; (5) Install the Docker Image in Flywheel (at this point, it is called a Flywheel Gear); (6) Run an analysis in Flywheel with the same test data used locally, to check that the results are the same.

#### Scitran: programmatic access to Flywheel

This is an important component of the whole RTP solution, as it allows programmatic control of the whole Flywheel system. In Figure 1 and Figure 2B we saw that the data storage and the computation jobs are always done in the server side. For a couple of subjects, the interaction with Flywheel can be done through the web interface and most of the settings and tools can be tested this way. But increasingly, neuroimaging projects consist of larger number of subjects, and programmatic access is required for time efficient and error free data, configuration parameter and result management. Even with two subjects, it is easy to make mistakes when setting the more than 50 RTP-pipeline config parameters.

With these objectives in mind, we developed the Scitran tool. Scitran is a Matlab application built on top of Flywheel’s Matlab SDK, that provides several functionalities: *(1) Data/analysis results search:* to perform any of the following operations, *(2) Metadata update:* for example, socioeconomic information from the subjects, *(3) Upload and download information:* log files, images, or other project level required information, *(4) Job launch and management:* select a gear, version, config parameters and launch an analysis job and control the status, *(5) Information extraction:* select the desired subjects and analysis and extract only the required results that go to the statistical analysis. Specifically for RTP-pipeline, there is a utility that extracts the per-subject-per-tract metric (e.g. FA) and creates a table with all the project level, acquisition level and analysis level parameters.

Scitran is publicly available and can be installed from (https://github.com/vistalab/scitran/). Installing it in Matlab is as simple as downloading the zip file or cloning the repository and adding it to the Matlab path.

Scitran comes with the latest Flywheel Matlab SDK. In order to make it work, it is required to have an active user in a Flywheel instance and downloading an API key. Please refer to the Scitran manual (https://github.com/vistalab/scitran/wiki) for further information on installing and using the Scitran tool.

### Data analysis pipeline

On top of the neuroinformatic platform and the programmatic access tool, we created the Docker Images with the DWI analysis pipeline that conform the total RTP solution. The Docker Images can be used without the platform, i.e. if the same Image is used and the same inputs and configuration parameters are provided, the results will be the same. The RTP solution is divided into two main, and some optional/auxiliary steps. The main steps are RTP-preproc and RTP-pipeline.

#### RTP-preproc

The preprocessing Gear prepares the data for the posterior DWI analysis in RTP-pipeline. The preprocessing does not include any data analysis, and its objective is to correct known problems in the DWI acquisitions process. It can optionally align the DWI data with the anatomical file. This gear is an adaptation from mrTrix’s (https://github.com/MRtrix3/mrtrix3) recommended preprocessing procedure implemented in Brainlife (https://brainlife.io/app/5a813e52dc4031003b8b36f9). In Figure 3A we see the required and optional inputs to the Gear:

- Required:

∘ DIFF. Diffusion data in 4D nifti format
∘ BVAL. Bvec file in FSL format
∘ BVEC. Bval file in FSL format
- Optional:

∘ ANAT. Anatomical T1w file: used to optionally align the diffusion data to it
∘ RDIF. Reversed phase encoding DWI data.
∘ RBVL. Bval corresponding to RDIF
∘ RBVC. Bvec corresponding to RDIF

**Figure 3.**
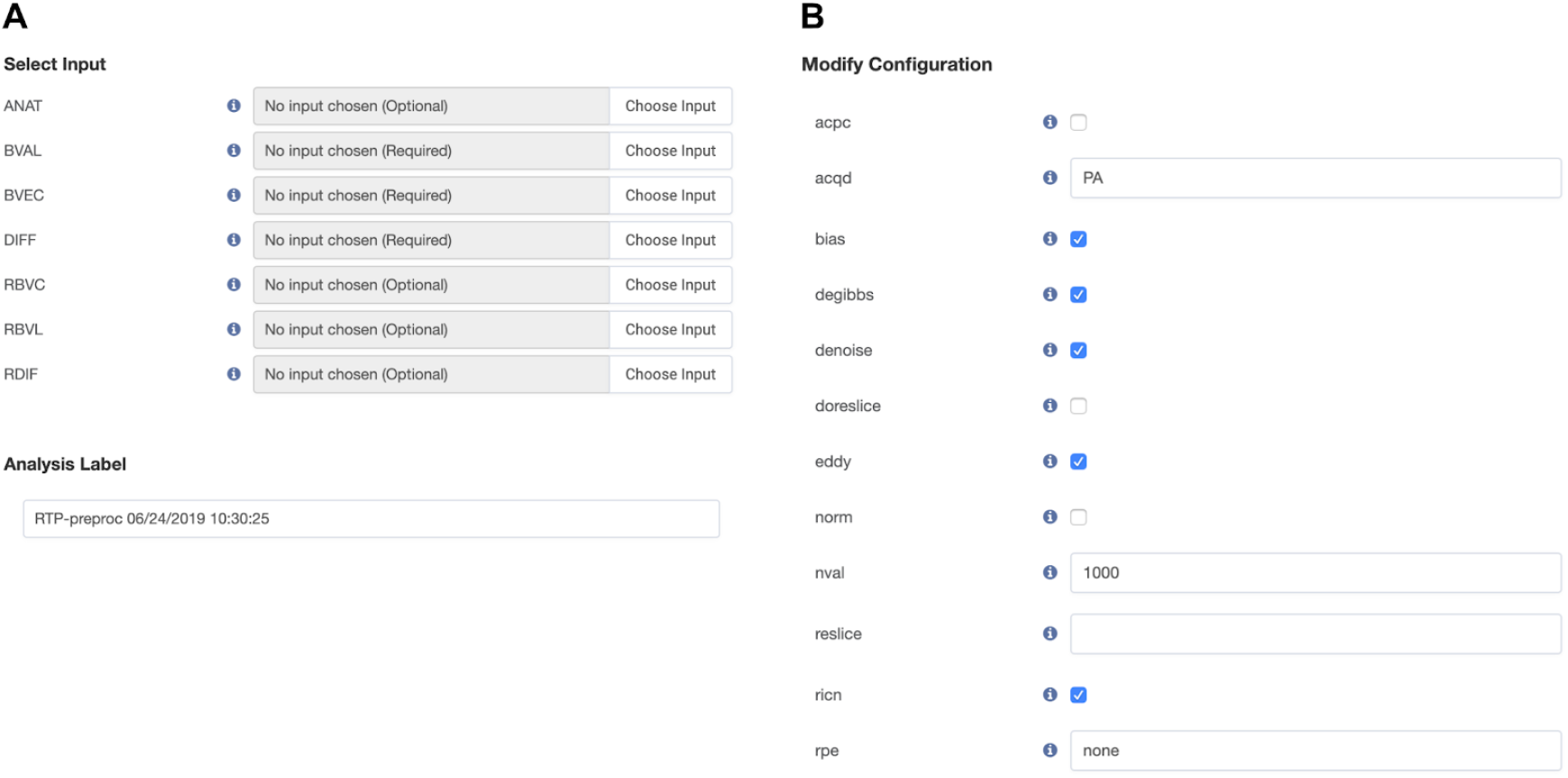
Input and parameter definition for RTP-preproc. (A) Required and Optional input files to the RTP-preproc gear. (B) List of configurable parameters for RTP-preproc, in alphabetical order (Flywheel’s default).

It is highly recommendable acquiring the reversed phase encoding files to correct for EPI geometric distortions with FSL’s TOPUP. If those images are not provided, the alignment between the diffusion data and the anatomy is not going to be as good. This can be relevant downstream for analysis that require a good alignment between anatomical and diffusion data (for example if we want to use mrTrix’s Anatomically Constrained Tractography).

Next we will provide an exhaustive list of the preprocessing steps that can be set using the configuration parameters seen in Figure 3B. The parameters in Figure 3B do not show a logical order, as Flywheel sorts them alphabetically. We will use an application logic order here, this is the chronological order how the options are applied:

1. rpe: specifies if there is reverse phase encoding data or not
2. acqd: acquisition phase encoding direction
3. denoise: perform a principal component analysis (PCA) based denoising of the data (Veraart et al. 2016; Veraart, Fieremans, and Novikov 2016)
4. degibbs: perform Gibbs ringing correction (Kellner et al. 2016)
5. eddy: perform FSL’s eddy current correction; if inverted phase encoding direction files are found, eddy will be done as part of FSL’s topup (Andersson and Sotiropoulos 2016)
6. bias: compute bias correction with ANTs (Tustison et al. 2010)
7. ricn: perform Rician background noise removal
8. norm: perform intensity normalization
9. nval: normalize the intensity of the FA white matter mask to this number
10. align: align dwi data with AC-PC anatomy
11. doreslice: do reslicing to the amount set in reslice
12. reslice: if doreslice, reslice the DWI data to an isotropic voxel of this value.

Once the analysis is run, there is the possibility to visualize the results directly in the web browser window, or download all or a selection of images locally for visual inspection. Additionally, we use to run another small Gear called RTP-bvec-flip-detect at this stage. RTP-bvec-flip-detect is a deep neural network solution to detect if there has been a flip in the sign of any of the directions in the bvec files and corrects them. Depending on the acquisition and preprocessing steps, one of the directions in the bvec file might have the sign changed. For example, the output from RTP-preproc comes with the x direction flipped (this is a known effect of using FSL). RTP-bvec-flip-detect will automatically correct the error and copy to the output folder of the analysis all the preprocessed data ready to be used as an input to RTP-pipeline. The code is publicly available in https://github.com/vistalab/RTP-bvec-flip-detect.

#### RTP-pipeline

The tractography container takes preprocessed DWI data and automatically identifies 22 white matter tracts (see Figure 4 for six illustrative tracts) and provides several diffusion profiles per each tract (see inset plot in Figure 4 for an illustrative example of two group mean FA tract profiles). In the analysis process, the pipeline generates values at the voxel and the tract level, that can be later used for further analysis. The required inputs to the Gear are the dwi, bvec and bval files, and can take an anatomical T1-weighted image and Freesurfer’s aparcaseg segmentation files as optional. Although the aparcaseg file is really optional and its usage depends on decisions downstream, we always encourage using the T1-weighted image for better results.

**Figure 4.**
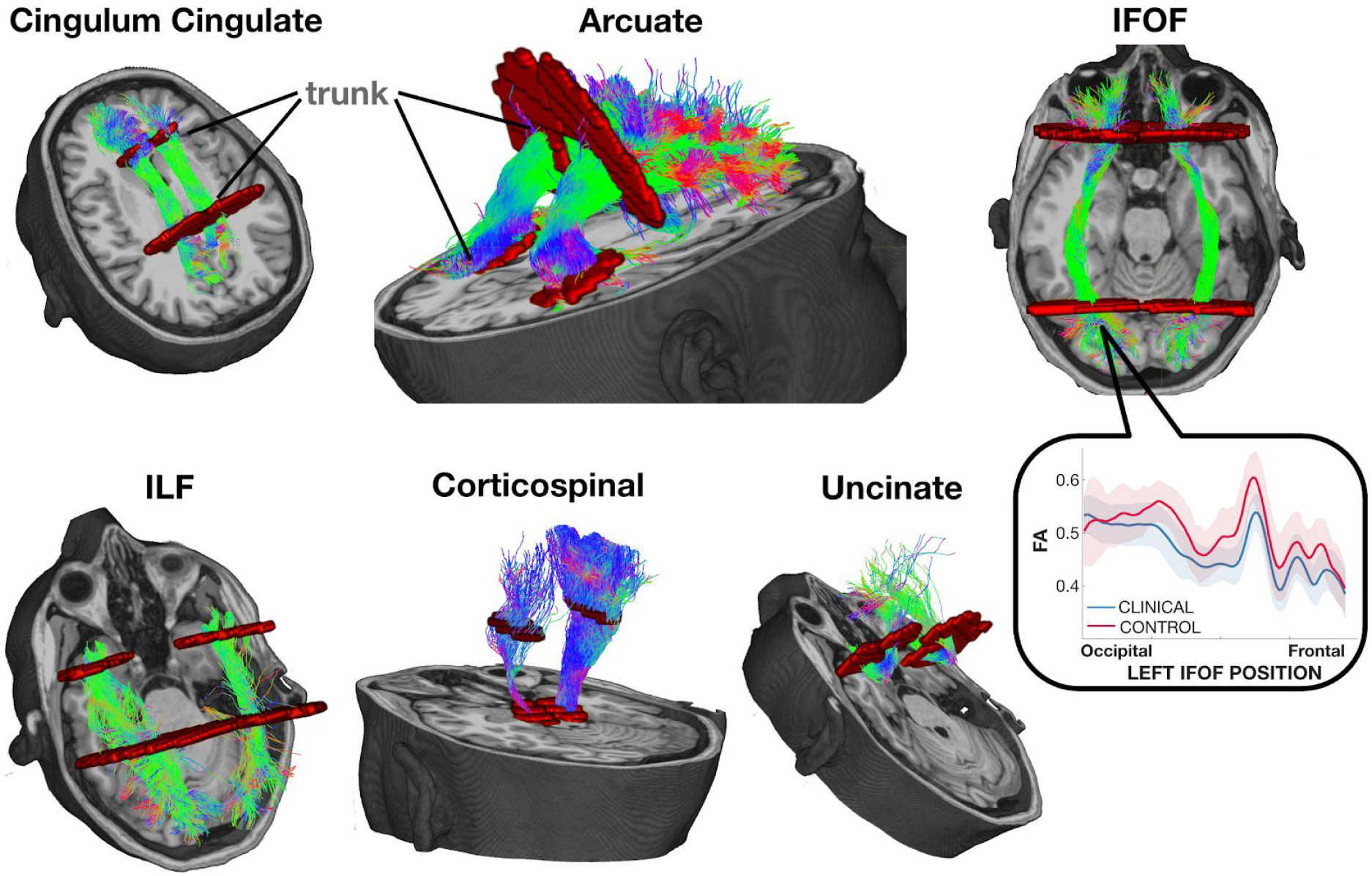
Six illustrative pairs of homologous tracts and their defining ROIs. The streamlines serve as a model of white matter tracts; they are selected by fitting to the diffusion weighted imaging (DWI) measurements. The tracts are defined by regions of interest (ROIs, red) that select specific streamlines from the whole brain tractogram. The region between the two ROIs is relatively stable and called the trunk. We estimate a core fiber from the collection of streamlines and sample 100 equally spaced segments. The FA of the core fiber is calculated by combining FA transverse to the core fiber at every sample point, using a Gaussian weighting scheme over distance. The set of sample points is the tract profile; the average of the FA values of the core fiber is the mean tract FA. In the inset, an illustrative example of two group profiles. The dark outline is the mean of the group and the shaded outline represents a standard deviation of the values.

In the current version, RTP-pipeline is heavily based on mrTrix, although legacy Matlab based code is still present in the code and can still be selected through configuration parameters (for example, a deterministic tractography implementation). The list of parameters is too long to show it here, for a complete list please refer to the manifest.json file in (github.com/vistalab/RTP-pipeline). The manifest.json contains a description and a default value per every parameter. There are many legacy parameters that will be removed in the future, but that are still in use/repurposed in the current version. It is important to notice that the parameter selection is project specific. We recommend to use the default in a first test run, but from that moment on, it is usually required to adjust the parameters. The github repository contains a wiki page with parameter recommendations (github.com/vistalab/RTP-pipeline/wiki/Parameter-recommendations) for different types of data, with the hope that it will be of help to other researchers. We hope that it will grow in the future. Here we will illustrate the main steps and will explain the main configuration parameter choices:

1. The diffusion data are aligned and resliced to the anatomical image (https://github.com/vistalab/vistasoft. dtiInit).
2. DTI modeling at the voxel level and metric creation: FA, MD, RD…
3. Response function creation. Different algorithms can be selected, and with different options. We recommend using the dhollander algorithm with automatic lmax, it separates the WM, GM, and CSF in the DWI without requiring an anatomical file.
4. Constrained Spherical Deconvolution (CSD) modelling. Different algorithms can be selected. Performed at the voxel level, it creates Fiber Orientation Distributions (FOD) that are required for tracking.
5. Whole brain white matter streamlines estimation. Different algorithms and options are available (see mrTrix documentation for a comprehensive list on the tckgen command). Additionally, there are other decisions that can be taken at this point. We implemented Ensemble Tractography (Takemura et al. 2016) and LiFE (Linear Fascicle Evaluation) (Pestilli et al. 2014) into the RTP-pipeline. Is is possible to run them independently, i.e. both, one or the other, or none can be selected.

a. Ensemble Tractography (ET) method: ET invokes MRtrix’s tractography tool a number of times (selected with a configuration parameter), constructing whole brain tractograms with a range of tracking parameter options. For example, the minimum angle parameters for tracking can be varied (e.g. 47, 23, 11, 6, 3), and the resulting whole brain connectomes concatenated together. There are config parameters specific to the Ensemble Tractography implementation.
b. The LiFE (Linear Fascicle Evaluation) method evaluates the tractogram streamlines and retains those that meaningfully contribute to predicting variance in the DWI data. There are config parameters specific to the LiFE implementation.
6. The Automated Fiber Quantification (AFQ) method (Yeatman et al. 2012) segments streamlines into tracts (Figure 4). The tracts are defined by regions of interest (ROIs, in red in Figure 4) that select specific streamlines from the whole brain tractogram. Several options that control this step can be configured with the configuration parameters (see the wiki for the most current options).
7. Tract Profiles. The voxel based metrics are calculated for each individual tract creating the tract profiles. The metrics can be of any voxel based measurement. Although the most common ones are the DTI metrics (FA, MD,…), we have successfully used others, such as T1 relaxation time or macromolecular tissue volume (MTV) quantitative MRI maps. The basic steps for tract profile creation are:

a. A core fiber, representing the central tendency of all the streamlines in the tract, is identified.
b. Equally spaced positions along the fiber between the two defining ROIs are sampled (N=100).
c. The metric values of streamlines at locations transverse to each sample position are measured and combined. The combination is weighted by a Gaussian value based on the distance from the sample point.
d. The sampling and transverse averaging generates a tract profile of 100 metric values (see inset in Figure 4 for a group mean FA tract profile).

### Data preparation and reproducible statistical analysis

Once the tract profiles have been generated, we can retrieve them for statistical analysis using Scitran methods. The data preparation, statistical analysis and plotting scripts read the data directly from the Flywheel neuroinformatics platform, selecting the values from the relevant analysis results. The Scitran tools include methods to download all the relevant data and metadata (acquisition metadata, gear version and configuration parameters, subject’s metadata…) together with the tract profiles, ready for analysis.

There are two ways to guarantee reproducibility of the statistical results:

1. The scripts are stored and versioned in a GitHub repository. The input data to those scripts is stored and versioned in the neuroinformatic platform. The scripts will always read the same data files, perform the same statistics and create the same plots. An example script that uses the Scitran tool to interact with Flywheel can be found at https://github.com/garikoitz/paper-reproducibility.
2. Once a paper is accepted for publication, a more permanent reproducible system can be implemented as a Gear that contains the scripts, all their dependencies and the manifest of all the computational parameter values. This Gear will be a snapshot in time of the statistical analysis scripts. It will read the data from Flywheel and produce the same results as in the published paper.

This last step guarantees the full cycle of computational reproducibility, from MRI data acquisition to result publication.

## Acknowledgements

This work was supported by a Marie Sklodowska-Curie (H2020-MSCA-IF-2017-795807-ReCiModel) grant to G.L.-U. We thank the Simons Foundation Autism Research Initiative and Weston Havens foundation for support.

## References

Andersson, Jesper L. R., and Stamatios N. Sotiropoulos. 2016. “An Integrated Approach to Correction for off-Resonance Effects and Subject Movement in Diffusion MR Imaging.” NeuroImage 125 (January): 1063–78.

Baker, Monya. 2016. “1,500 Scientists Lift the Lid on Reproducibility.” Nature 533 (7604): 452–54.

Buckheit, Jonathan B., and David L. Donoho. 1995. “WaveLab and Reproducible Research.” Wavelets and Statistics. https://doi.org/10.1007/978-1-4612-2544-7_5.

Carp, Joshua. 2012. “On the Plurality of (methodological) Worlds: Estimating the Analytic Flexibility of FMRI Experiments.” Frontiers in Neuroscience 6 (October): 149.

Kellner, Elias, Bibek Dhital, Valerij G. Kiselev, and Marco Reisert. 2016. “Gibbs-Ringing Artifact Removal Based on Local Subvoxel-Shifts.” Magnetic Resonance in Medicine: Official Journal of the Society of Magnetic Resonance in Medicine / Society of Magnetic Resonance in Medicine 76 (5): 1574–81.

Marcus, Daniel, John Harwell, Timothy Olsen, Michael Hodge, Matthew Glasser, Fred Prior, Mark Jenkinson, Timothy Laumann, Sandra Curtiss, and David Van Essen. 2011. “Informatics and Data Mining Tools and Strategies for the Human Connectome Project.” Frontiers in Neuroinformatics 5: 4.

Peng, Roger D. 2011. “Reproducible Research in Computational Science.” Science 334 (6060): 1226–27.

Pestilli, Franco, Jason D. Yeatman, Ariel Rokem, Kendrick N. Kay, and Brian A. Wandell. 2014. “Evaluation and Statistical Inference for Human Connectomes.” Nature Methods 11 (10): 1058–63.

Sandve, Geir Kjetil, Anton Nekrutenko, James Taylor, and Eivind Hovig. 2013. “Ten Simple Rules for Reproducible Computational Research.” PLoS Computational Biology 9 (10): e1003285.

Stodden, Victoria, Friedrich Leisch, and Roger D. Peng. 2014. Implementing Reproducible Research. CRC Press.

Takemura, Hiromasa, Cesar F. Caiafa, Brian A. Wandell, and Franco Pestilli. 2016. “Ensemble Tractography.” PLoS Computational Biology 12 (2): 1–22.

Tustison, Nicholas J., Brian B. Avants, Philip A. Cook, Yuanjie Zheng, Alexander Egan, Paul A. Yushkevich, and James C. Gee. 2010. “N4ITK: Improved N3 Bias Correction.” IEEE Transactions on Medical Imaging 29 (6): 1310–20.

Veraart, Jelle, Els Fieremans, and Dmitry S. Novikov. 2016. “Diffusion MRI Noise Mapping Using Random Matrix Theory.” Magnetic Resonance in Medicine: Official Journal of the Society of Magnetic Resonance in Medicine / Society of Magnetic Resonance in Medicine 76 (5): 1582–93.

Veraart, Jelle, Dmitry S. Novikov, Daan Christiaens, Benjamin Ades-Aron, Jan Sijbers, and Els Fieremans. 2016. “Denoising of Diffusion MRI Using Random Matrix Theory.” NeuroImage 142 (November): 394–406.

Wilson, Greg, Jennifer Bryan, Karen Cranston, Justin Kitzes, Lex Nederbragt, and Tracy K. Teal. 2017. “Good Enough Practices in Scientific Computing.” PLoS Computational Biology 13 (6): e1005510.

Yarkoni, Tal, and Jacob Westfall. 2017. “Choosing Prediction Over Explanation in Psychology: Lessons From Machine Learning.” Perspectives on Psychological Science: A Journal of the Association for Psychological Science 12 (6): 1100–1122.

Yeatman, Jason D., Robert F. Dougherty, Nathaniel J. Myall, Brian A. Wandell, and Heidi M. Feldman. 2012. “Tract Profiles of White Matter Properties: Automating Fiber-Tract Quantification.” PloS One 7 (11): e49790.

